# Proteomic analysis reveals microvesicles containing NAMPT as mediators of radiation resistance in glioma

**DOI:** 10.1101/2022.03.23.485479

**Authors:** Elena Panizza, Brandon D. Regalado, Fangyu Wang, Ichiro Nakano, Nathaniel M. Vacanti, Richard A. Cerione, Marc A. Antonyak

**Affiliations:** Department of Molecular Medicine, Cornell University, Ithaca, NY 14853; Tsukuba and Ehime Universities, Japan; Division of Nutritional Sciences, Cornell University, Ithaca, NY 14853; Department of Chemistry and Chemical Biology, Cornell University, Ithaca, NY 14853

**Keywords:** brain cancer, glioma, glioblastoma, proteomics, radio-resistance, radiation, systems biology, NAD^+^, NAMPT, extracellular vesicles, cellular proliferation, therapy resistance

## Abstract

Glioma is a malignant brain tumor that is highly resistant to radiation and chemotherapy, where patients survive on average only 15 months after diagnosis. Furthering the understanding of mechanisms leading to radiation resistance of glioma is paramount to identify novel therapeutic targets. Previous studies have shown that glioma stem cells (GSCs) play an important role in promoting radiation resistance and disease recurrence. Herein we analyze the proteomic alterations occurring in patient-derived GSCs upon radiation treatment in order to identify molecular drivers of resistance. We show that proteome changes upon radiation accurately predict the resistance status of the cells, whereas resistance to radiation does not correlate with glioma transcriptional subtypes. We further show that the radio-resistant GSC-267 cell line sheds microvesicles (MVs) enriched in the metabolic enzyme nicotinamide phosphoribosyltransferase (NAMPT). These MVs can be transferred to recipient fibroblasts and radio-sensitive GSCs, enhancing their intracellular total NAD^+^ and NADH level, and their ability to proliferate when cultured in low serum, treated with a radio-mimetic drug or irradiated. The NAMPT enzymatic inhibitor FK-866 blocked the ability of MVs from GSC-267 cells to mediate these effects. Similarly, GSC-267 cells where NAMPT was knocked-down using shRNA, which produced MVs depleted of this enzyme, were unable to promote cell proliferation. Collectively, our findings demonstrates that proteome-level regulation can accurately predict the radio-resistance status of GSCs, and identifies NAMPT transfer via MVs as a mechanism for spreading radiation resistance within the glioma tumor microenvironment.

**Significance:** The highly aggressive and deadly brain cancer glioma is commonly resistant to standard chemo- and radio-therapy. We used systems biology approaches to study patient-derived glioma stem cells (GSCs), which are known to be responsible for therapeutic resistance, and cell-to-cell communication mediated by extracellular vesicles (EVs), which plays an important role in tumor progression. Analysis of the proteome of GSCs and of the EVs they release led us to determine that the EV-mediated transfer of the metabolic enzyme nicotinamide phosphorybosyltransferase (NAMPT) from radio-resistant to less aggressive cells confers resistance to radiation. Our findings identify a mechanism of therapy resistance in glioma, and suggest that NAMPT inhibition could enhance the efficacy of radiation for the treatment of glioma.

## Introduction

Glioma is an aggressive brain cancer with poor clinical outcomes. The disease is typically treated with surgical resection followed by chemotherapy and radiotherapy. However, the tumor invariably reappears, progresses rapidly, and is resistant to treatment. Over the last 15 years, many research efforts have been undertaken to further our understanding of the mechanisms that promote glioma, with the hope that they will lead to the development of new therapeutic strategies. In 2006, the existence of glioma stem cells (GSCs) was first reported (1). Shortly thereafter, the genomic landscape of glioma was described based on large cohorts of patients (2), facilitating the identification of key genetic and epigenetic alterations, i.e. the discovery of mutations in isocitrate dehydrogenase 1 and 2 (IDH1 and IDH2) and the glioma-CpG island methylator phenotype (g-CIMP), which are associated with a relatively favorable prognosis (3, 4). In addition, molecular subtyping of glioma was established based on transcriptional signatures (5). Despite these advances in understanding the disease, only two approaches (temozolomide and tumor treating fields) have been approved by the U.S. Food and Drug Administration for the treatment of newly diagnosed glioma in over two decades (6, 7). A wider range of options exist for treatment of recurrent disease (8), including the only targeted approach approved for glioma, i.e. the drug bevacizumab, an antibody that inhibits vascular endothelial growth factor (VEGF) (9). However, glioma remains a major challenge confronting oncologists, with most patients surviving only 12-18 months following their initial diagnosis (10).

Radiation therapy kills cancer cells by producing extensive damage to their DNA. GSCs have generated a good deal of interest after the discovery that they play a causative role in chemotherapy- and radio-resistance (1, 11, 12). A major contributor to drug and radiation resistance are extracellular vesicles (EVs) produced and shed by GSCs. There are two major classes of EVs, microvesicles (MVs) that bud off the plasma membrane and exosomes which form from intraluminal vesicles within multivesicular bodies. EVs produced by aggressive cancer cells, including GSCs, can be transferred to less aggressive or lower-grade cancer cells and potentiate their proliferation, therapy resistance and invasiveness. Cancer cell-derived EVs have also been shown to play an important role in communicating with the surrounding stroma and shaping the tumor microenvironment (13–15). Yet, there are still significant gaps in our understanding of the mechanisms by which EVs mediate their effects, particularly as they relate to GSCs.

In the current study, we quantified the proteomic alterations occurring in several patient-derived GSC lines following their irradiation. The analysis led us to identify a subset of GSC lines that are resistant to radiation and another subset that is radio-sensitive. Intriguingly, we found that MVs, but not exosomes, produced by the radio-resistant GSC-267 cell line promote the proliferation of both fibroblasts and radio-sensitive GSCs cultured in low serum, irradiated, or treated with the radiation mimetic Bleomycin. Proteomic analysis of the content of the EVs produced by GSC-267 cells led to the identification of the metabolic enzyme nicotinamide phosphoribosyltransferase (NAMPT) as a key cargo of MVs. NAMPT is responsible for regenerating nicotinamide adenine dinucleotide (NAD^+^) from nicotinamide as part of the NAD^+^ salvage pathway (16, 17). It has been reported to promote the progression of several types of cancers (18–21). One way NAMPT contributes to the cancer process is by providing NAD^+^ that serves as a co-factor for sirtuin-1 (SIRT1), which in turns deacetylates the tumor suppressor p53 on lysine residue 382, resulting in p53 inhibition and suppression of apoptosis (22–24). NAMPT also increases the activity of poly(ADP-ribose) polymerases (PARPs), another class of NAD^+^-consuming enzymes, leading to enhanced resistance to DNA damage (21). Here we show that GSC-267 cells in which NAMPT was knocked-down, shed MVs that are depleted of this enzyme which are incapable of promoting the proliferation of irradiated cells. These findings highlight how GSCs are able to communicate with their surroundings by shedding a specific class of EVs to promote resistance to radiation. They also raise the interesting possibility that strategies to block the production of MVs, or inhibit the activity of NAMPT, can potentially be combined with radiation to more effectively treat glioma.

## Results

### Proteomic profiling identifies a subset of GSC lines that are resistant to radiation

GSCs are thought to be a major source of therapy resistance in glioma (25–28). To gain a better understanding of the mechanisms used by GSCs to promote radiation resistance, liquid chromatography tandem mass spectrometry (LC-MS/MS) was used to analyze the proteome of eight GSC lines, each derived from an individual patient. GSCs were either left untreated, or were treated with six Gray (Gy) of ionizing radiation, one tenth of the full therapeutic dose that is typically administered to patients over six weeks (29). To obtain the relative quantification of proteins in the samples, peptides were labeled using tandem mass tags (TMT) 10-plex. The peptides were then fractionated using high-resolution isoelectric focusing (HiRIEF) to facilitate the identification of an elevated number of peptides, providing comprehensive coverage of the proteome and high quantitative accuracy (30–32). Overall, 120,883 unique peptides corresponding to 9,108 proteins that map to unique genes were identified (**SI Appendix, Fig. S1A**).

The examined cell lines are representative of different glioma molecular subtypes, as defined by Verhaak and colleagues (5, 33) (**SI Appendix, Fig. S1C**). To determine how each GSC line responds to radiation at the proteomic level, individual protein abundances were normalized to the average of untreated and treated for each cell line (**SI Appendix, Table S1**; see **Materials and Methods**). Hierarchical clustering based on Euclidian distance and Ward method (34) separates two subsets of GSC lines within treatment groups (**Fig. 1A**). In one subset, the expression levels of 2,658 proteins were found to change in response to radiation. This group is referred to as the radio-sensitive group. In the other subset, the expression level of only 86 proteins changed, and these GSC lines are considered as the radio-resistant group (**Fig. 1B**). To substantiate the classification as radio-resistant and radio-sensitive cell lines, proliferation assays were performed on several of the GSC lines after they were irradiated. **Figure 1C** shows that the proliferation of radio-sensitive GSC-408 and GSC-374 cell lines (green) is significantly diminished after radiation treatment, while the proliferation of radio-resistant GSC-267 and GSC-84 cell lines (red) is not impacted. Additionally, radio-sensitive GSCs display significant changes in the expression of proteins that are well-known transcriptional targets of p53, which orchestrates a series of cellular responses to DNA damage (35). These include the upregulation of p21 (CDKN1A), which is responsible for cell cycle arrest upon DNA damage (36); ribonucleotide-diphosphate reductase subunit M2 B (RRM2B), which is necessary for DNA synthesis during DNA repair (37); tumor necrosis factor receptor superfamily member 6 (FAS), which mediates p53-dependent apoptosis after DNA damage (38); and the downregulation of protein aurora borealis (BORA), which is required for progression through mitosis (39) (**Fig. 1D**). However, changes in the levels of the same proteins were not observed in radio-resistant GSCs upon treatment with radiation. At this point, we wanted to determine whether there is an association between the GSCs radio-sensitivity status and known molecular signatures that are used to describe glioma. Three of the radio-sensitive GSC lines (GSC-157, GSC-408 and GSC-1079) were found to be IDH mutant or hypermethylated (g-CIMP +) based on an established gene expression signature (40, 41). The remaining two radio-sensitive (GSC-374 and GSC-1037) and three radio-resistant GSC lines were determined to be IDH wild type (**SI Appendix, Fig. S1B**). Additionally, among the radio-resistant GSC lines identified in our analysis, only GSC-267 cells display a mesenchymal (MES) signature, which has been associated with a more aggressive phenotype (42) (**SI Appendix, Fig. S1C**). This suggests that transcriptional subtyping and IDH mutational status do not fully describe resistance to radiation, in line with the concept that additional markers may be necessary to better predict clinical outcomes of glioma (43–46).

**Figure 1.**
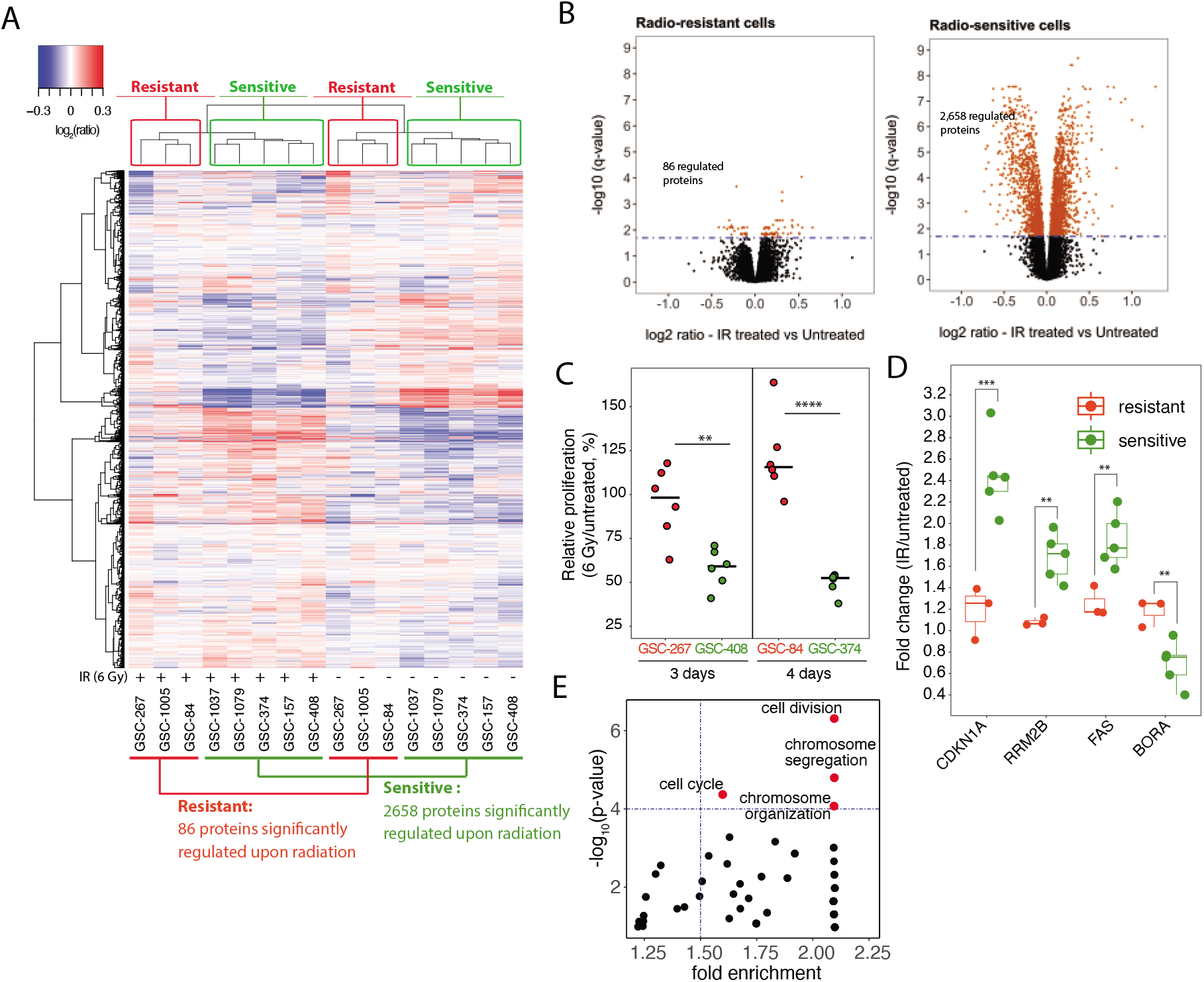
Proteomic analysis identifies a subset of GSC lines that are resistant to radiation. A. A subset of GSC lines that do not respond to radiation is separated by hierarchical clustering based on Euclidian distance and Ward linkage method. The number of proteins significantly regulated in radio-resistant (red) and radio-sensitive (green) GSCs upon radiation treatment is displayed. IR, ionizing radiation. B. Volcano plots representing changes in protein expression upon radiation, plotted against the significance of the regulation. Each dot represents the average log2(IR treated/untreated) for each protein in the subset of radio-resistant (left) or radio-sensitive (right) GSC lines. Proteins whose expression is significantly changed are selected by using the statistical package limma with a Bonferroni-Hochberg corrected q-value lower than 0.02. C. Relative proliferation of representative radio-resistant (red) and radio-sensitive (green) GSCs following their treatment with 6 Gy of ionizing radiation. Dots represent independent biological replicates. Significance levels were evaluated using Student’s t-test. D. Box plot representing the change in expression of the indicated proteins upon radiation treatment. Significance levels were evaluated using Student’s t-test. E. GO biological processes enriched in the subset of proteins elevated in radio-resistant versus radio-sensitive GSC lines. Proteins elevated in radio-resistant compared with radio-sensitive GSC lines were selected using the statistical package limma with a Bonferroni-Hochberg corrected q-value lower than 0.02 and a log2(ratio) > 0 (n= 81). The terms “cell division”, “cell cycle”, “chromosome segregation” and “organization” are significantly enriched (red dots), based on Fisher’s exact test p-value lower than 1*10^−4^ and fold enrichment higher than 1.5. In all figure panels, significance of observed changes is represented as: **, p-value < 1*10^−2^; ***, p-value < 1*10^−3^.

Since the expression of a relatively small number of proteins change in the radio-resistant subtype of GSCs upon irradiation, we hypothesize that these cells have undergone alterations that make them less sensitive to radiation. To pinpoint alterations that are characteristic of radio-resistant GSCs, we examined their steady-state protein abundances (**SI Appendix, Table S2**). We found an enrichment of the cell cycle, cell division and chromosome segregation processes when considering proteins that are elevated in radio-resistant compared with radio-sensitive GSCs (**Fig. 1E**). This suggests that radio-resistant GSCs potentially overcome the consequences of radiation-induced DNA damage by taking advantage of an increased ability to progress through cell cycle and proliferate.

### Microvesicles derived from the radio-resistant GSC-267 cell line are enriched in NAMPT and promote proliferation

EVs are known to be important for the ability of GSCs to communicate with their surrounding and promote different aspects of cancer progression. Thus, we were interested in determining whether EVs plays an important role in mediating therapy resistance. We observed that the radio-resistant GSC-267 cell line displays an enrichment of the gene ontology (GO) term “vesicle-mediated transport” compared with the other cell lines (**SI Appendix, Fig. S2A**, **Fig. 2A**). Vesicle-mediated transport includes the processes of vesicle formation, coating, budding and fusion with target membranes, and has been shown to describe both MVs and exosomes in a previous proteome-wide analysis (47). We then assayed how EVs released by GSC-267 cells would affect the proliferation of fibroblasts deprived of nutrients. NIH/3T3 cells cultured in 0.5% calf serum (CS, low serum) were either left untreated or were treated with MVs or exosomes isolated from GSC-267 cells. While exosomes had little effect, MVs significantly increased the proliferation of recipient cells (**Fig. 2B**).

**Figure 2.**
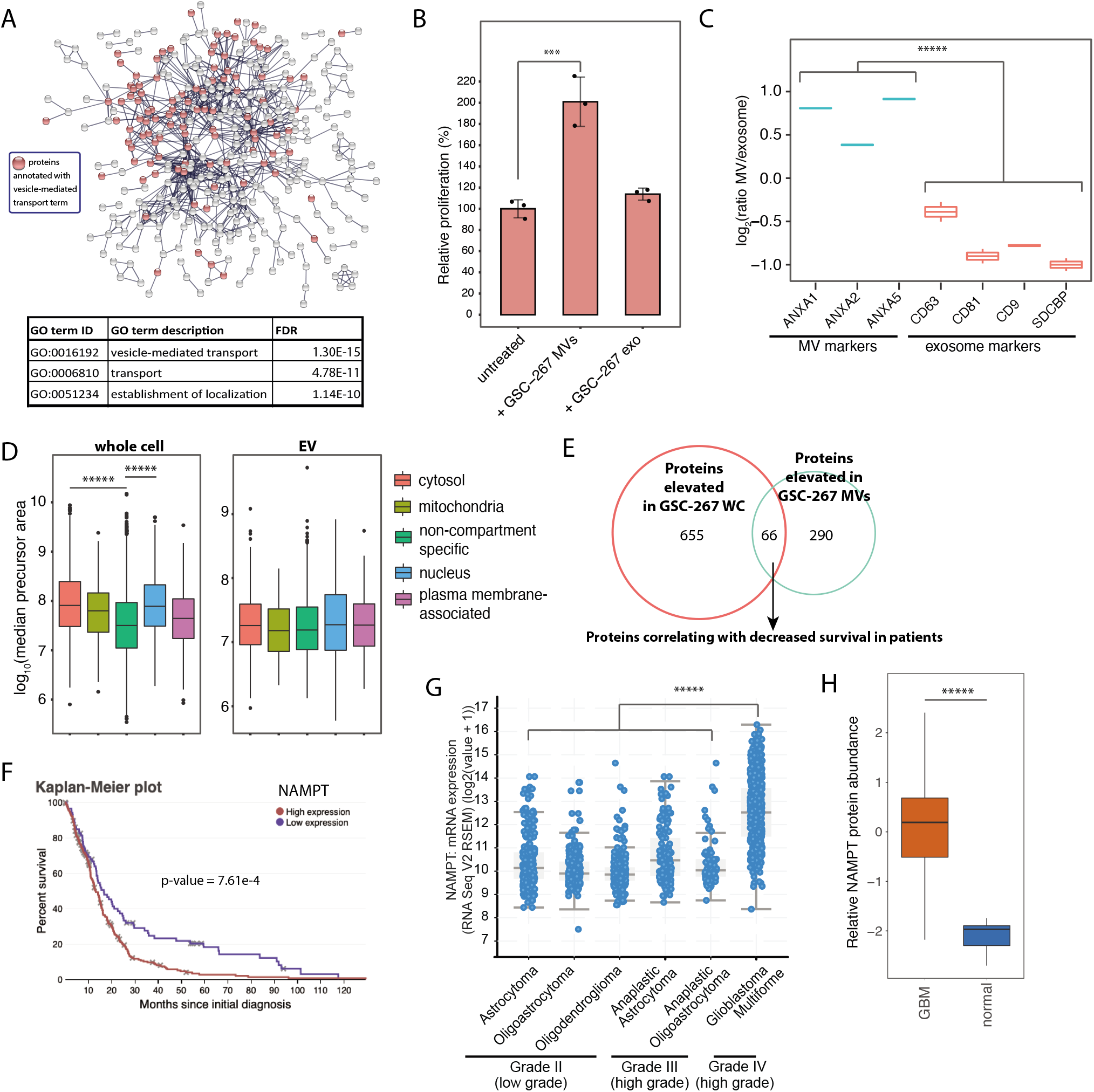
Microvesicles derived from GSC-267 cells increase the proliferation of recipient cells and are enriched in NAMPT. A. Proteins elevated in GSC-267 cells, compared with other GSC lines, are highly connected in a protein-protein interaction network (protein-protein interaction enrichment p-value < 1*10^−16^). The top 3 GO terms enriched in the network are listed. FDR, false discovery rate-adjusted p-value for the enrichment based on Fisher’s exact test. B. Relative proliferation of NIH/3T3 cultured in low serum (0.5% calf serum, CS), either untreated or treated with MVs or exosomes isolated from GSC-267 for 5 days. The displayed p-value was calculated using the Student’s t-test. Individual dots represent independent biological replicates. C. Protein expression levels of EV markers in MVs relative to exosomes isolated from GSC-267 cells (n = 2 for each sample type). ANXA1, 2, 5, Annexin A1, A2, A5; SDCBP, Syntenin-1. The displayed p-value was calculated using the Student’s t-test. D. Distribution of protein abundances based on precursor areas in whole cell (left panel) and EV (right panel) proteomic analyses. Proteins are categorized by subcellular compartment (49). The number of compartment-enriched proteins in each category is > 300 in the whole cell analysis and >= 70 in the EV analysis. The displayed p-value was calculated using the Student’s t-test. E. Identification of proteins that are candidates for promoting the ability of MVs derived from GSC-267 cells to confer aggressive phenotypes. Proteins that are specifically elevated in GSC-267 whole cell (WC) lysates compared with the other GSC lines (n=721), and proteins that are elevated in MVs generated by GSC-267 cells compared with exosomes (n=356) are displayed; the overlap between these two sets (n=66) was selected as candidates. This set of proteins was then filtered based on correlation with decreased survival in a TCGA patient cohort (n = 349). F. Elevated NAMPT transcript expression significantly correlates with lower survival of patients diagnosed with glioblastoma (GBM, grade IV glioma) (n= 349). The displayed p-value was calculated using a logrank test based on data obtained from TCGA. The 25^th^ percentile value of NAMPT expression values is set as a threshold to define high expression. G. NAMPT transcript expression levels across GBM (grade IV), high-grade glioma (grade III) and low-grade glioma (grade II). The displayed p-value was calculated using a Student’s t-test based on a TCGA cohort of 1,520 patients. H. Relative NAMPT protein abundance in glioblastoma samples (n=99) compared with normal brain tissue (n=10). The displayed p-value was calculated using the Student’s t-test. In all figure panels, significance of observed changes is represented as: ***, p-value < 1*10^−3^; *****, p-value < 1*10^−6^.

To better understand how MVs mediate this phenotype, we compared the proteomic content of MVs and exosomes generated by GSC-267 cells using LC-MS/MS. Two replicates were analyzed for each sample, resulting in the identification of 1,252 proteins (**Fig. S2B, SI Appendix, Table S3**). MV markers, including annexin 1, 2, and 5 (ANXA1, ANXA2, ANXA5), are elevated in the MV samples, while the exosome markers CD81, CD9 and syntenin-1 (SDCBP) (48) are increased in the exosome samples (**Fig. 2C**). Moreover, the expression of cytosolic and nuclear proteins (49) is significantly elevated in whole-cell samples, but not in EV samples, as compared to non-compartment-specific proteins (**Fig. 2D**). For example, the well-established cytosolic and nuclear markers GAPDH, HISTH1B, RPS3, LMNA and PSMA3 (49) have a much higher relative abundance in whole-cell compared with EV samples (**SI Appendix, Fig. S2C**). This analysis demonstrates that GSC-267 cells are capable of generating both MVs and exosomes and that the EV fractions isolated from these cells are devoid of cellular contaminants.

A set of 66 proteins are elevated in both GSC-267 cells, and in the MVs that they produce, but not in their exosomes (**SI Appendix, Fig. S2D, Fig. 2E**). These proteins were considered as potential candidates for mediating the proliferation-promoting properties of the MVs. Within this set of proteins, a further selection based on their correlation with decreased patient survival highlighted two proteins (**SI Appendix, Fig. S3A**), including the metabolic enzyme nicotinamide phosphoribosyltransferase (NAMPT), which has been heavily implicated in cancer progression (18–21). NAMPT is the rate-limiting enzyme in the NAD^+^ salvage pathway, responsible for the production of NAD^+^ from nicotinamide. Several studies have shown that the ability of NAMPT to generate NAD^+^ is important in promoting glioma as well as other cancers. Elevated NAMPT transcript expression correlates with poor glioma patient survival (**Fig. 2F**), and NAMPT transcript is significantly elevated in grade IV (glioblastoma, GBM) compared with lower grade glioma (**Fig. 2G**). Furthermore NAMPT protein expression is also significantly elevated in GBM samples compared with normal brain tissue based on a large proteomic study of glioma (**Fig. 2H**) (45).

### The ability of MVs from GSC-267 cells to promote radio-resistance is dependent on NAMPT enzymatic activity

Our results indicated that NAMPT is elevated in radio-resistant GSC-267 cells and within the MVs that they generate. GSC-267 cells have the highest expression level of NAMPT in the panel (**Fig. 3A**). The radio-sensitive GSC lines identified in our analysis all have lower levels of NAMPT, and the MVs derived from these cells lack the enzyme. Interestingly, we did find an example of a radio-resistant cell line, GSC-84, that expresses relatively high levels of NAMPT; however, their MVs do not contain the enzyme (**Fig. 3B**). Therefore, we compared the ability of MVs derived from GSC-84 cells (which lack NAMPT) and MVs from GSC-267 cells (which do contain the enzyme), to enhance the proliferation of NIH/3T3 cells treated with Bleomycin, a drug that mimics the effects of radiation. Bleomycin treatment strongly decreased fibroblast proliferation, an effect that could be overcome by treating the cells with MVs isolated from GSC-267 cells. In contrast, MVs isolated from GSC-84 cells failed to restore cell proliferation (**Fig. 3C**), suggesting that the presence of NAMPT within MVs may be important for promoting radio-resistance. We next examined whether MVs derived from GSC-267 cells could similarly promote the proliferation of irradiated GSC-1079 cells, a radio-sensitive cell line that expresses some of the lowest levels of NAMPT (**Fig. 3A**). The results in **Fig. 3D** show that these MVs can indeed rescue GSC-1079 cells from the inhibitory effect of ionizing radiation on cell proliferation.

**Figure 3.**
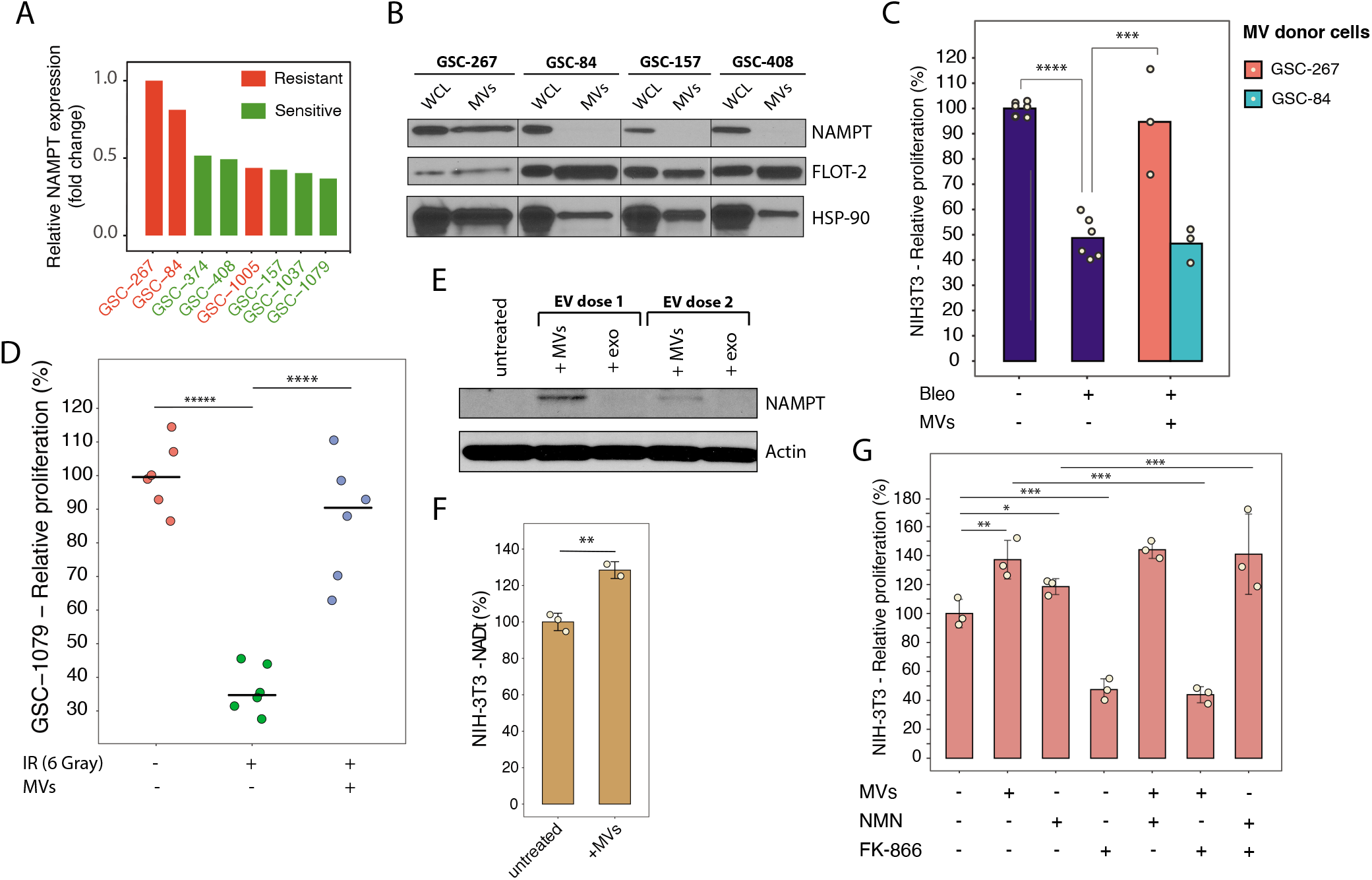
NAMPT enzymatic activity is required to promote the proliferation of MV-recipient cells. A. Relative protein expression levels of NAMPT across GSC lines based on proteomic analysis. B. Western blot representing the expression levels of NAMPT and the MV markers flotillin-2 (FLOT-2) and heat shock protein 90 (HSP-90) across GSC lines and in the MVs derived from these cells. WCL, whole cell lysate. C. Relative proliferation of NIH/3T3 cells cultured in low serum (0.5% CS) medium, either left untreated or treated with the indicated combinations of 5 μM bleomycin (Bleo) and MVs derived from GSC-267 or GSC-84 cells for 5 days. Individual dots represent independent biological replicates. D. Relative proliferation of radio-sensitive GSC-1079 cells either untreated or treated with 6 Gy of ionizing radiation (IR) alone or in combination with MVs derived from GSC-267 cells for 4 days. Individual dots represent independent biological replicates. E. Western blot representing the expression levels of NAMPT in NIH/3T3 cells, either untreated or treated with progressively lower doses of MVs and exosomes (exo) isolated from GSC-267 cells (EV dose 2 is half of EV dose 1) for 24 hours. F. Total NAD^+^ and NADH level (NADt) in NIH/3T3 cells, untreated or treated with MVs derived from GSC-267 cells for 6 hours. Individual dots represent independent biological replicates. G. Relative proliferation of NIH/3T3 cultured in low serum (0.5% CS), either untreated or treated with MVs derived from GSC-267 cells, 500 μM nicotinamide mononucleotide (NMN), and 10 μM of the NAMPT inhibitor FK-866, as indicated, for 3 days. Individual dots represent independent biological replicates. In all figure panels, significance of observed changes was evaluated using Student’s t-test. *, p-value < 0.02; **, p-value < 1*10^−2^; ***, p-value < 1*10^−3^; ****, p-value < 1*10^−4^.

NAMPT has been shown to promote cancer progression through its enzymatic activity which increases intracellular NAD^+^ levels (21), as well as by acting as a cytokine-like molecule upon its secretion from cells (50). Recent studies have demonstrated that secreted NAMPT (extracellular NAMPT, eNAMPT) can act as a ligand for the cell surface receptor C-C chemokine receptor type 5 (CCR5) (51, 52). Binding of eNAMPT to CCR5 leads to enhanced proliferation of muscle stem cells, while CCR5 hyperactivation has been linked with breast cancer progression (53). Therefore, we considered whether NAMPT was expressed along the surfaces of MVs from GSC-267 cells (54, 55), such that it would be able to engage and activate CCR5 in recipient cells. However, blocking CCR5 with its antagonist cenicriviroc did not inhibit the ability of MVs to rescue the proliferation of NIH/3T3 cells upon Bleomycin treatment (**SI Appendix, Fig. S3B**). Moreover, the addition of human recombinant NAMPT, which can bind and activate CCR5 (52, 56), failed to rescue the proliferation of irradiated fibroblasts (**SI Appendix, Fig. S3C**).

To determine whether the enzymatic activity of NAMPT is necessary for the ability of MVs produced by GSC-267 cells to promote the proliferation of stressed cells, we examined whether NAMPT protein is transferred to recipient cells via MVs. **Fig. 3E** shows that human NAMPT is present in lysates from NIH/3T3 mouse fibroblasts upon treatment with MVs, and that the amount of enzyme detected in the cells is proportional to the dose of MVs used for treatment. Fibroblasts treated with MVs derived from GSC-267 cells also have a ~30% increase of the total intracellular NAD^+^ and NADH level (NADt), compared to untreated fibroblasts (**Fig. 3F**). Nicotinamide mononucleotide (NMN) is the product of NAMPT and can be used to mimic the effects of the active enzyme. The addition of NMN and MVs from GSC-267 cells increased the proliferation of fibroblasts that were cultured in low serum to a similar extent as treatment with MVs alone. Moreover, treatment with the NAMPT enzymatic inhibitor FK-866 prevented the MV-mediated increase in proliferation, but did not affect the proliferation of fibroblasts that were also treated with NMN (**Fig. 3G**). Collectively, these findings demonstrate that the ability of MVs derived from GSC-267 cells to rescue the proliferation of recipient cells exposed to radio-mimetic treatment, low serum and radiation treatment is dependent on the enzymatic activity of NAMPT that the MVs transfer.

### NAMPT transfer via MVs confers resistance to radiation onto recipient cells

To determine whether NAMPT within MVs is necessary to confer radio-resistance, a GSC-267 cell line was generated that has stably integrated a construct encoding for a small hairpin RNA (shRNA) targeting NAMPT. The expression of the construct is controlled by doxycycline (Dox), so that upon addition of Dox the expression of NAMPT is knocked-down in a time-dependent manner (**SI Appendix, Fig. S4D**). This cell line, referred to as GSC-267 NAMPTsh, was supplemented with NMN to support normal cell metabolic activity and ability to generate MVs. The MVs derived from the control (-Dox) GSC-267 NAMPTsh cells contained NAMPT, while the MVs from the cells induced to express the NAMPT-targeting shRNA (+Dox), had less of the enzyme (**Fig. 4A-B**). GSC-267 NAMPTsh +Dox cells were able to generate similar numbers of EVs as cells expressing NAMPT (-Dox), based on nanoparticle tracking analysis (NTA) (**Fig. 4C-D**).

**Figure 4.**
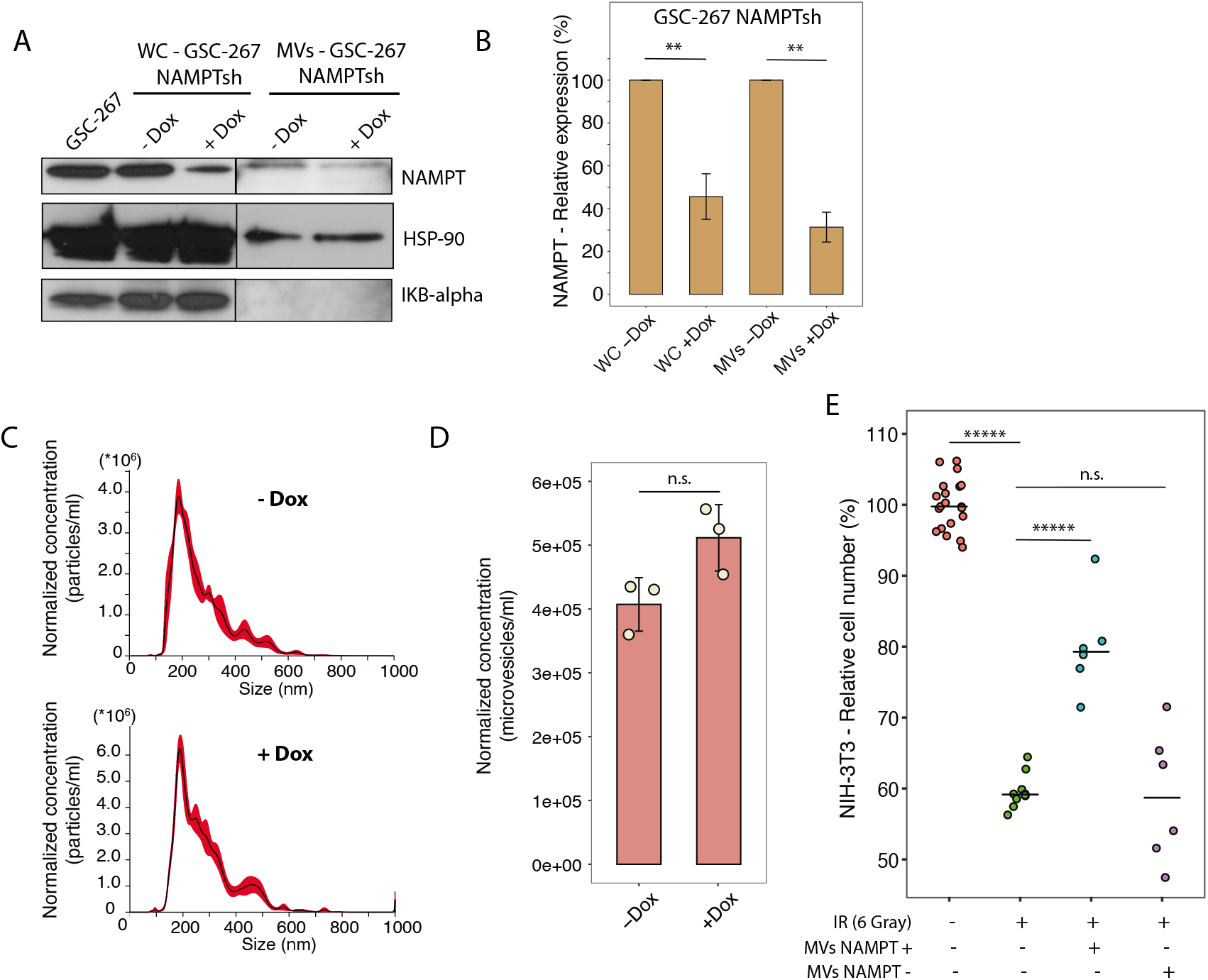
The MV-mediated transfer of NAMPT to cells confers resistance to ionizing radiation. A. NAMPT protein expression levels in whole cell (WC) lysates from GSC-267 parental cells, GSC-267 NAMPTsh cells −/+ Dox, and in MV protein extracts derived from GSC-267 NAMPTsh cells −/+ Dox. HSP-90 protein expression level was used as loading control. IKB-alpha is a WC marker and shows that the MV preparations were devoid of cellular contaminants. B. Densitometric quantification of NAMPT protein expression level as displayed in 4B. Variability is evaluated based on two technical replicates. C. Representative images of nanoparticle tracking analysis (NTA) of the partially clarified medium (containing EVs) collected from GSC-267 NAMPTsh cells-Dox (top panel) or +Dox (bottom panel). Exosomes (vesicles smaller than 200 nm) and MVs (vesicles larger than 200 nm) are present. D. Concentration of MVs (vesicles larger than 200 nm) isolated from GSC-267 NAMPTsh cells −/+Dox, normalized based on cell number. Individual dots represent independent biological replicates. E. Relative number of viable NIH/3T3 cells cultured in low serum (0.5% CS). Cells were either left untreated or treated with 6 Gy of ionizing radiation (IR) alone or in combination with MVs derived from GSC-267 NAMPTsh cells - Dox (MVs NAMPT +), or + Dox (MVs NAMPT-), for 4 days. Individual dots represent independent biological replicates. In all figure panels, significance of observed changes was evaluated using Student’s t-test. n.s., not significant; **, p-value < 1*10^−2^; *****, p-value < 1*10^−6^.

Finally, we treated irradiated NIH/3T3 cells with equal numbers of MVs derived from either the NAMPT expressing (-Dox), or NAMPT knockdown (+Dox), GSC-267 NAMPTsh cells, and then assayed their proliferation. While the MVs containing NAMPT significantly enhanced the proliferation of irradiated cells, the MVs depleted of NAMPT failed to rescue cell proliferation (**Fig. 4E**). These findings demonstrate that the transfer of NAMPT to recipient cells mediated by MVs promotes radiation resistance.

## Discussion

This study presents a previously undescribed mechanism for radio-resistance in GBM that involves the MV-mediated transfer of the metabolic enzyme NAMPT to recipient cells. The expression of NAMPT is essential for normal cell metabolism (17), since it regulates the intracellular NAD^+^ pool by catalyzing the first step in the biosynthesis of NAD^+^ from nicotinamide. NAMPT has been shown to restore NAD^+^ levels and counteract the effects of ageing (57, 58). NAMPT is also overexpressed in several different pathological contexts, such as obesity, liver disease, diabetes, and cancer. Indeed, a number of lines of evidence point to NAMPT playing an important part in the development of aggressive cancers (18, 59, 60). It has been shown to be highly expressed in tumor tissues from large cohorts of cancer patients, and to correlate with decreased patient survival (**Fig. 2F-G; refs**). Historically, some of the effects of NAMPT have been attributed to a freely secreted form of the enzyme, referred to as eNAMPT, which can act as a ligand for cell surface receptors (17). However, it is becoming increasingly clearer that intercellular communication mediated by EVs plays important roles in mediating NAMPT functions. EVs containing NAMPT have been shown to promote the proliferation of muscle stem cells, in manner that is dependent on NAMPT enzymatic activity (52). A recent report also showed that NAMPT-containing EVs can reverse the effects of ageing in mice (56). A small number of studies have identified NAMPT as a cargo protein within EVs isolated from the plasma of melanoma and breast cancer patients (61, 62), as well as being present within EVs shed by breast cancer cells (63). Here we demonstrate that NAMPT is enriched in MVs produced by radio-resistant GSCs (**Fig. 3B, SI Appendix, Fig. S3A**), and that its transfer to recipient cells can restore their proliferative capability upon radiation treatment (**Fig 3C-D, Fig. 4E**). Further studies will be needed to clarify the specific roles of EV-contained NAMPT for cancer progression.

Cultures of GSCs derived from individual patients reliably recapitulate the cellular diversity of the original tumors (42). GSCs are considered critical drivers of resistance to radiation and chemotherapy (1, 11). Previous reports have identified targets that could sensitize GSCs to radiation (1, 25, 64, 65); however, so far none have progressed to the clinic. Here we present a comprehensive proteomic profile of a set of patient-derived GSC lines in response to radiation (**Fig. 1A-B**). We identified a subset of radio-sensitive GSC lines (**Fig. 1A-C**) that stop proliferating upon irradiation and display a strong molecular response to DNA damage, including changes in the expression of proteins that direct cell cycle arrest, DNA repair and apoptosis (**Fig. 1D**). We also discovered a second subset of radio-resistant GSC lines which do not display the typical molecular response to DNA damage caused by radiation, and instead continue to proliferate (**Fig. 1A-D**). Two of the radio-resistant GSC lines overexpress NAMPT (**Fig. 3A**); however, only one of these cell lines, GSC-267, sheds MVs that contain the enzyme (**Fig. 2A, SI Appendix, Fig. S3A, Fig. 3B**). This suggests that the three radio-resistant GSC lines are representative of tumors with distinct mechanisms of resistance to radiation, and a future goal will be to define them. The three radio-resistant GSC lines identified in this analysis belong to different transcriptional subtypes as defined by Verhaak and colleagues (5). Recently, the proteomic landscape of GBM has been described by the Clinical Proteomic Tumor Analysis Consortium (CPTAC), based on a collection of 99 treatment-naïve tumor samples (45). In the CPTAC study, proteome-level information captured a subset of patients that belong to a different subtype compared with previously defined transcriptional subtypes. Additionally, in recent years, studies that employ proteomics-based molecular profiling and other large-scale methods (66) to describe the clinical landscape of several types of cancer have been published. Novel insights into immune-based subtyping, oncogenic drivers and patient stratification are only few among the discoveries that these studies have brought about, and which could not be captured by genomic or transcriptomic analyses alone (31, 45, 67–70). In this study we demonstrate that proteome-level changes occurring upon treatment can accurately describe glioma radio-sensitivity status. This finding confirms that proteome-level information can lead to important new knowledge, as well as highlights the importance of continuing to study brain tumors at the proteome-level.

The characterization and definition of EVs is an active field of research and many recent advances have been made (71–73). A relatively small number of studies has used systems biology approaches that broadly characterize the transcriptomic and proteomic content of EVs, and most studies still rely on the detection of a few specific markers (74). This study presents a comprehensive overview of the proteome of EVs generated by GSCs. The analysis identified over 1,200 proteins contained within EVs (**SI Appendix, Fig. S2B, Table S3**). The large number of proteins detected allowed for unbiased target discovery (**Fig. 2E**), and the comprehensive proteome of GSC-derived EVs presented here can be used as a resource for future studies (**SI Appendix, Table S1 and S2**). Well-established EV markers could be reliably detected within MVs and exosomes using proteomics (**Fig. 2D**). Loss of enrichment in intracellular proteins was used to effectively demonstrate the lack of cellular contaminants in EV preps (**Fig. 2D** and **SI Appendix, Fig. S2C**), thus demonstrating that systems biology approaches can be applied to determine the purity and identity of EV preparations.

Thus far, targeting NAMPT with small molecule inhibitors in clinical trials has resulted in toxicity (75–80). However, the apparent important role played by this enzyme in cancer progression warrants the examination of alternative strategies that impact its regulation. For example, NAMPT overexpression is known to contribute to radiation resistance by increasing intracellular NAD^+^ levels, which in turn facilitate DNA damage repair via the PARP family of enzymes. Elevated NAD^+^ levels can also protect cells from radiation-induced DNA damage by activating autophagy (64, 81) (82, 83). Therefore, it may prove worthwhile to target the PARP enzymes or to inhibit autophagy as a strategy for treating GBM cases where NAMPT is highly expressed. Moreover, because NAMPT has been detected directly in plasma and cerebrospinal fluid samples (84–88), both in pathological (cancer and other diseases) and in physiological contexts, it might also serve as a valuable marker for prognosis or for selecting patients for therapy.

## Materials and Methods

### Cell culture and treatment

GSC lines GSC-267, GSC-1005, GSC-84, GSC-1037, GSC-1079, GSC-374, GSC-157 and GSC-408 were established from freshly resected glioma tumor samples in the laboratory of Dr. Nakano as previously described (42, 65, 89). Cells were cultured in Dulbecco’s Modified Eagle Medium:Nutrient Mixture F-12 (DMEM/F-12, Gibco) with the addition of B-27 supplement to a final concentration of 2% v/v (Gibco), heparin (2.5 mg/ml, Sigma-Aldrich), basic fibroblast growth factor (bFGF, 20 ng/ml, Pepro Tech. Inc), epidermal growth factor (EGF, 20 ng/ml, Pepro Tech. Inc) and penicillin and streptomycin to a final concentration of 100 IU/ml and 100 μg/ml respectively (both from Sigma-Aldrich). GSC spheres were dissociated mechanically by pipetting or using TrypLE Express Enzyme (ThermoFisher), and bFGF and EGF were added twice a week. For proteomic analysis, GSCs were seeded in 175 cm^2^ flasks using fresh complete medium at a confluency of about 50-60% and a volume of 50 ml two days before radiation treatment. GSCs were either left untreated or administered a single dose of 6 Gray (Gy) of ionizing radiation using a Mark I Model 68 Cesium-137 gamma irradiator at the Irradiator Facility within the Transgenic Mouse Core Facility at Cornell University. Cells were harvested for proteomic analysis 48 hours after irradiation. NIH/3T3 cells (ATCC CRL-1658) were cultured in DMEM media (Gibco) supplemented with 10% (v/v) CS (Gibco) and penicillin and streptomycin to a final concentration of 100 IU/ml and 100 μg/ml respectively. Compounds used for treatment include the following: bleomycin sulfate (Cayman Chemicals, cat. no. 13877); cenicriviroc, (Selleck Chemicals, cat. no. S8512); β-Nicotinamide mononucleotide, ≥95% HPLC (Sigma Aldrich, Millipore Sigma, cat. no. N3501); Recombinant Human Visfatin (PeproTech, cat. no. 130-09); FK-866 (Cayman Chemical Company, cat. no. 13287). When EVs were used to treat cells, equal amounts were added to each experimental well. EVs derived from 125,000 EV donor cells were used to treat individual wells containing 2,000 recipient cells and were added every 24 hours.

### Protein extraction

GSCs were transferred to conical tubes and pelleted by centrifugation at 100 x *g*. The cells were washed twice with Phosphate-Buffered Saline (PBS, Corning) and dry cell pellets were stored at −80°C. Cells were lysed in the presence of 4% (w/v) sodium dodecyl sulfate (SDS), 25 mM HEPES pH 7.6 and 1 mM dithiothreitol (DTT, Sigma) (4% SDS cell lysis buffer). Adherent cells were scraped off the culture plates in the presence of 4% SDS cell lysis buffer. Lysates were then heated at 95°C for 10 min, sonicated and then centrifuged for 15 min at 14,000 x *g* and 4°C. Supernatants were transferred to new vials, and protein concentrations were quantified using the DC Protein Assay (Bio-Rad).

### Protein digestion for whole cell proteomic analyses

Protein extracts were processed following the Single-Pot Solid-Phase-enhanced Sample Preparation (SP3) protocol (90, 91) with slight modifications. Briefly, a bead slurry was prepared by mixing 50 μl of Sera-Mag SpeedBeads Carboxyl Magnetic Beads (hydrophobic) with 50 μl of Sera-Mag SpeedBeads Carboxyl Magnetic Beads (hydrophilic) (both from GE Healthcare). The bead slurry was washed two times with 200 μl of MQ water using a magnetic rack, prior to resuspension in 500 μl of MQ water. Protein samples (200 μg each) were diluted to a final volume of 200 μl using the cell lysis buffer described above. Cysteine residues were alkylated by adding 40 μl of SP3 bead slurry to the protein samples, and chloroacetamide (Sigma) to a final concentration of 40 mM, and acetonitrile to a final concentration of 70% (to bind proteins to the SP3 beads). Samples were then incubated for 20 minutes on a rotating rack. Following the incubation, supernatants were removed and SP3 beads were washed twice with 200 μl of 70% EtOH and once with 180 μl of 100% acetonitrile using a magnetic rack. SP3 beads were air-dried for 30 seconds prior to digestion at 37°C overnight in 100 μl of a solution containing 50 mM HEPES pH 7.6, 1 M urea and 4 ug of Lysyl Endopeptidase (Lys-C, Wako) with mild shaking. Subsequently, 100 μl of a solution containing 50 mM HEPES pH 7.6 and 4 ug trypsin (Pierce) was added and samples were incubated again at 37°C overnight with mild shaking. Supernatants were transferred to new tubes and peptide concentrations were quantified using the DC Protein Assay (Bio-Rad). 50 μg of each sample were set aside for TMT labeling. Identical linker samples were prepared to function as denominator in each TMT set. Linker samples were prepared by pooling equal amounts of proteins from each of the 16 samples, up to a final amount of 50 μg each. All samples were lyophilized prior to TMT labeling.

### TMT labelling

Peptide samples were labeled with 10-plex TMT reagents (Thermo Scientific) as previously described in Panizza *et al.* (30). Briefly, before labeling samples were resuspend using TEAB pH 8.5 (50 mM final concentration) to adjust the pH. Each sample was labeled with an isobaric TMT-tag. Labelling efficiency was determined by LC-MS/MS before pooling the samples. Pooled samples were desalted with Reversed Phase-Solid Phase Extraction cartridges (Phenomenex, Torrance, CA, USA) and then lyophilized in a SpeedVac (Thermo Fisher Scientific) Sample clean-up was performed by solid phase extraction (SPE strata-X-C, Phenomenex). Purified samples were dried in a SpeedVac.

### Peptide-level HiRIEF

High-resolution isoelectric focusing (HiRIEF) was performed as previously described in Branca et al. (32). Briefly, peptides from whole cell samples were focused on immobilized pH gradient (IPG) gel strips (GE Healthcare, Waukesha, WI, USA) on a linear pH range of 3-10. Strips were divided into 72 fractions and extracted to V-bottom 96-well plates with a liquid handling robot (GE Healthcare prototype modified from Gilson liquid handler 215). Plates were lyophilized in a SpeedVac prior to LC-MS/MS analysis. Dried peptides were dissolved in 3% ACN/0.1% FA and consolidated into 40 fractions based on fraction complexity. Specifically, less complex fractions are pooled: i.e., fractions 19-26, 31-35, 42-49, 53-63, 66-70.

### EV isolation

EVs were isolated as described by Wang et al (74). Briefly, medium containing GSCs was subjected to two consecutive centrifugations at 100 × g to clarify the medium of cells and debris. The partially clarified medium was filtered using a 0.22 μm pore size Steriflip PVDF filter (Millipore). MVs were collected off the filter by adding 4% SDS cell lysis buffer for protein extraction, or cell culture medium for functional assays. The filtrate was subjected to ultracentrifugation at 100,000 × g for 3 h to collect exosomes. The exosome pellet was either collected by adding 4% SDS cell lysis buffer for protein extraction, or cell culture medium for functional assays.

### Protein extraction, digestion and TMT labeling for EV proteomic analysis

To quantify the proteome of EVs derived from the GSC-267 cell line, cells were seeded in seven 175 cm^2^ flasks using fresh complete medium at a confluency of about 60-70% and a volume of 50 ml the day before EV collection. MVs and exosomes were isolated as detailed above, and proteins were extracted using 130 μl of 4% SDS lysis buffer. Protein digestion was performed using the SP3 protocol described above for whole cell samples, but volumes were adapted as follows. EV protein samples (~10-40 μg each) were diluted to a final volume of 120 μl using 4% SDS cell lysis buffer. Cysteine residues were alkylated by adding chloroacetamide (Sigma) to a final concentration of 40 mM, 6 μl of SP3 bead slurry, and acetonitrile to a final concentration of 70% (to bind proteins to the SP3 beads). Samples were then incubated for 20 minutes on a rotating rack. Following the incubation, supernatants were removed and SP3 beads were washed twice with 100 μl of 70% EtOH and once with 100 μl of 100% acetonitrile using a magnetic rack. SP3 beads were air-dried for 30 seconds prior to digestion at 37°C overnight in 20 μl of a solution containing 50 mM HEPES pH 7.6, 1 M urea and 0.6 ug of Lysyl Endopeptidase (Lys-C, Wako) with mild shaking. Subsequently, 20 μl of a solution containing 50 mM HEPES pH 7.6 and 0.6 ug trypsin (Pierce) was added and samples were incubated again at 37°C overnight with mild shaking. Supernatants were transferred to new tubes and peptide concentrations were quantified using the DC Protein Assay (Bio-Rad). Peptides were labeled using 10-plex TMT reagents as described above and 4 μg of proteins were used for each TMT channel. Linker samples were prepared by pooling four EV samples, corresponding to a final amount of 4 μg of peptides.

### LC-MS/MS

LC-MS/MS analysis was performed by the Clinical Proteomics Mass Spectrometry facility at Karolinska Institutet-Karolinska University Hospital, Science for Life Laboratory in Stockholm, Sweden. Peptide samples were separated using a reversed-phase gradient containing phase A solution (5% DMSO, 0.1% FA) and phase B solution (90% acetonitrile, 5% DMSO, 5% water, and 0.1% formic acid). For whole cell proteomic analysis, each HiRIEF fraction was analyzed independently using a gradient that proceeded from 6% phase B to 37% phase B over 30-90 min depending on the complexity of the fraction being analyzed. For EV proteomic analysis, peptides were separated using a gradient that proceeded from 6% phase B to 30% phase B over 180 min. Upon completion of the gradient, the column was washed with a solution of 99% phase B for 10 min and re-equilibrated to the initial composition. A nano Easy-Spray column (PepMap Rapid Separation Liquid Chromatography; C18; 2-μm bead size; 100Å pore size; 75-μm internal diameter; 50 cm long; Thermo Fisher Scientific) was used on the nanoelectrospray ionization Easy-Spray source at 60°C. Online LC-MS/MS was performed using a hybrid Q Exactive mass spectrometer (Thermo Fisher Scientific). Fourier transform–based mass spectrometer (FTMS) master scans with a resolution of 60,000 (and mass range 300-1500 m/z) were followed by data-dependent MS/MS (35,000 resolution) on the 5 most abundant ions using higher-energy collision dissociation (HCD) at 30% normalized collision energy. Precursor ions were isolated with a 2 m/z window. Automatic gain control targets were 1 × 10^6^ for MS1 and 1 × 10^5^ for MS2. Maximum injection times were 100 ms for MS1 and 100 ms for MS2. The entire duty cycle lasted ~1.5 s. Automated precursor-ion dynamic exclusion was used with a 60 s duration. Precursor ions with unassigned charge states or a charge state of +1 were excluded. An underfill ratio of 1% was applied.

### Proteomics database search and ratio calculation

Raw MS/MS files for the whole cell proteomics analysis were converted to mzML format using msConvert from the ProteoWizard tool suite (v3.0.19127) (92). Spectra were then searched using MSGF+ (v2018-07-21) and Percolator (v3.01) (93) using the Galaxy platform (94). The reference database was the *Homo sapiens* protein subset of Swissprot, canonical isoforms, released on 2018-08-02. MSGF+ settings included precursor ion mass tolerance of 10 ppm and peptide spectral matches (PSMs) allowed for up to two missed trypsin cleavages. Carbamidomethylation on cysteine and TMT 10-plex on lysine and the N terminus were set as fixed modifications, and oxidation of methionine was set as a dynamic modification while searching all MS/MS spectra. Quantification of TMT 10-plex reporter ions was performed using OpenMS project’s IsobaricAnalyzer (v2.0) (95). A false discovery rate cutoff of 1% was applied at the PSM level. To obtain relative quantification of steady-state protein abundances for whole cell samples, ratios were first normalized to the median of each TMT channel, assuming equal peptide loading of all samples. Then, the TMT reporter ion value for each PSM was divided by the TMT reporter ion value for channel 131 (the linker) in each TMT set, in order to normalize protein quantification across the two TMT sets. Finally, the median ratio of PSMs belonging to a unique gene symbol was used to obtain gene-centric protein quantifications. TMT ratios were then log2 normalized (**SI Appendix, Table S2**). To obtain relative quantification of protein level changes upon radiation, protein TMT ratios were subtracted with the average of untreated and treated for each individual cell line and each protein. This normalization method allows to highlight protein expression changes upon treatment (**SI Appendix, Table S1**). MS/MS spectra for the EV proteomic analysis were matched against the human and bovine subset of the Swissprot database, release 2018-08-2, using Sequest/Percolator under the Proteome Discoverer software platform (PD 1.4, Thermo Scientific). Settings for the search were the same as specified in Panizza et al. (30). Peptides that matched to the bovine reference proteome filtered out. To calculate TMT ratios, first the TMT reporter ion value for each PSM was divided by the TMT reporter ion value for channel 128C (the linker in set 1), or 130C (the linker in set 2), in order to normalize protein quantification across the two TMT sets. Then, TMT ratios were log2 normalized and the average of the four EV samples (two MV, two exosome samples) for GSC-267 cells was subtracted to each ratio for each individual protein to obtain relative protein abundances in MV compared with exosomes (**SI Appendix, Table S3**). The mass spectrometry proteomics data have been deposited to the ProteomeXchange Consortium via the PRIDE partner repository (https://www.ebi.ac.uk/pride/archive/) with the dataset identifier PXD030092.

### cGIMP/IDH mutational status and transcriptional subtyping

cGIMP+/IDH mutant status was determined based on an established 100 gene signature (40, 41). Transcriptional subtypes for the five IDH wild type GSC lines were determined based on gene signatures published by Verhaak and colleagues (5). For both analyses, gene set enrichment analysis (GSEA) scores were calculated using the R package GSVA (96) specifying “ssgsea” as method. To assign samples to a mutational status or to a subtype, a threshold score was defined based on GSEA scores for a set of 99 GBM samples (45) with a known mutational status and molecular subtype.

### Data visualization and bioinformatic analyses

All plots were generated using RStudio. The statistical package linear models for microarray data (limma) (97) was used to define significantly regulated proteins for all proteomic data analyses. GO enrichment analysis was performed using the R package TopGO (98) and Fisher’s exact test was employed to evaluate the significance of the enrichment. Protein-protein interaction analysis was performed using the web tool provided by the Search Tool for the Retrieval of Interacting Genes/Proteins (STRING; https://string-db.org) (99). High confidence interactions (interaction score > 0.900) were considered for the analysis. Label-free quantification was performed as described before (100). Briefly, protein MS1 precursor areas were calculated as the average of the top three most intense peptides for each TMT set. Then, the median of the two TMT sets was used as MS1 precursor area. Inferred gene identity false discovery rates were calculated using the picked-FDR method. Patient survival was analyzed in relation to NAMPT transcript expression based on data generated by TCGA using the Affymetrix platform HT HG U133A (2, 101), through the web tool available at https://www.betastasis.com.

### Cell proliferation and counting assay

Relative cellular proliferation was determined using the Cell Counting Kit-8 (CCK-8) assay (Dojindo Molecular Technologies, Inc., cat. no. CK04), following the manufacturer’s instruction. Briefly, cells were seeded in 96-well plates and treated for the indicated amount of time before adding a volume of 10 μl of CCK-8 reagent. The reaction was incubated for 2-6 hours before reading absorbance using a plate reader. The absorbance values of cell culture wells that contained only cell culture medium was used as a background and subtracted from the absorbance values obtained from the experimental wells. Relative proliferation was calculated as a percentage as indicated. Number of cells was counted using an imaging cytometer (Celigo, Nexcelom Bioscience) as previously described (102). Briefly, cells were stained using Hoechst dye (Life Technologies) to determine total cell number, and propidium iodide (Thermo Fisher) to determine the number of dead cells. The number of live cells is reported and was obtained by subtracting dead cell from total cell number.

### Measurement of NADt

Intracellular total NAD^+^ and NADH levels were measured using the NAD/NADH-Glo assay (Promega Corporation) following supplier instructions. Briefly, cells were seeded in white, flat-bottom 96-well plates (Costar) the day before the experiment. Cells were treated for 6 hours with MVs or left untreated. Cells were then incubated with 50 μl of NAD/NADH Glo detection reagent. After 1 h incubation, luminescence was determined in a microplate reader.

### Western blotting

For western blot analysis, equal amounts of proteins were separated using Novex WedgeWell 4-20%, Tris-Glycine gels (Invitrogen, Thermo Fisher Scientific). Proteins were transferred onto PVDF membranes (Thermo Fisher). Membranes were blocked with a solution of 5% non-fat dry milk (Bio-Rad Laboratories) in Tris Buffered Saline −0.5% Tween-20 (TBS-T), then incubated with primary antibody in a solution of 5% bovine serum albumin (BSA) (Bio-Rad Laboratories) in TBS-T overnight at 4 °C. Membranes were then washed in TBS-T solution, and incubated with horseradish peroxidase (HRP)-conjugated secondary antibody (anti-mouse, #7076, or anti-rabbit, #7074, Cell Signalling Technology, Inc.) in 5% non-fat dry milk in TBS-T, for 1 h at room temperature. After additional washes, membranes were incubated with a chemiluminescent detection reagent (Western Lightning Plus, Chemiluminescent Substrate, PerkinElmer Inc.), and imaged on HyBlot CL Autoradiography Film (Thomas Scientific), and the film was developed using developer and fixer solutions (Merry X-Ray Imaging, Inc.). The following primary antibodies were employed: HSP90 (#4877), IκBα (#4812), NAMPT (#6122), β-actin (#3700) and Flotillin-2 (#3436), all from Cell Signaling Technology, Inc. Quantification of Western blots was performed using the ImageJ software (103).

### Generation of GSC-267 NAMPTsh cells

A shRNA targeting NAMPT (sequence: GCTAGCAGCGATAGCTATGACATTTATTACT-AGTATAAATGTCATAGCTATCGCTTTTTT) was selected using the Genetic Perturbation Platform, from the Broad Institute (https://www.broadinstitute.org/genetic-perturbation-platform). The shRNA oligo was obtained as a duplexed DNA from Integrated DNA Technologies, Inc. and cloned into the EZ-Tet-pLKO-Puro vector (AddGene) using the InFusion ligation kit (Takara Bio). The cloned vector was sequenced to verify appropriate insertion of the shRNA. Lentiviruses were generated by transfecting HEK-293T cells with the shRNA plasmid, and the packaging plasmids (#12259 and #12263, AddGene) using Polyethylenimine (Thermo Fisher). The viruses shed into the medium by the cells were harvested 24 and 48 h after transfection. GSC-267 cells were infected by treatment with the virus and polybrene (8 μg/mL).

### NanoSight analysis

The size and concentration of EVs were determined using a NanoSight NS300 (Malvern, Cornell NanoScale Science and Technology Facility) as described previously (104). Briefly, GSCs were grown in absence of the B-27 supplement for 24 hours, before conditioned medium was collected and centrifuged twice at 100 *x g* for 5 minutes to pellet cells and debris. The partially clarified medium was then diluted in PBS and injected into the beam path to capture movies of EVs as points of diffracted light moving rapidly under Brownian motion. Five 45-second videos of each sample were taken and analyzed to determine the concentration and size of the individual EVs based on their movement, and then results were averaged together.

## Acknowledgements

This work was supported by the National Center for Research Resources (grant no. S10RR023781, providing the Mark I Model 68 Cesium-137 gamma irradiator) and by NIH grant CA201402. We also thank the Cornell Stem Cell and Transgenic Core Facility and especially Robert Munroe for their assistance with irradiating GSCs.

---

**Supplementary Figure S1.**
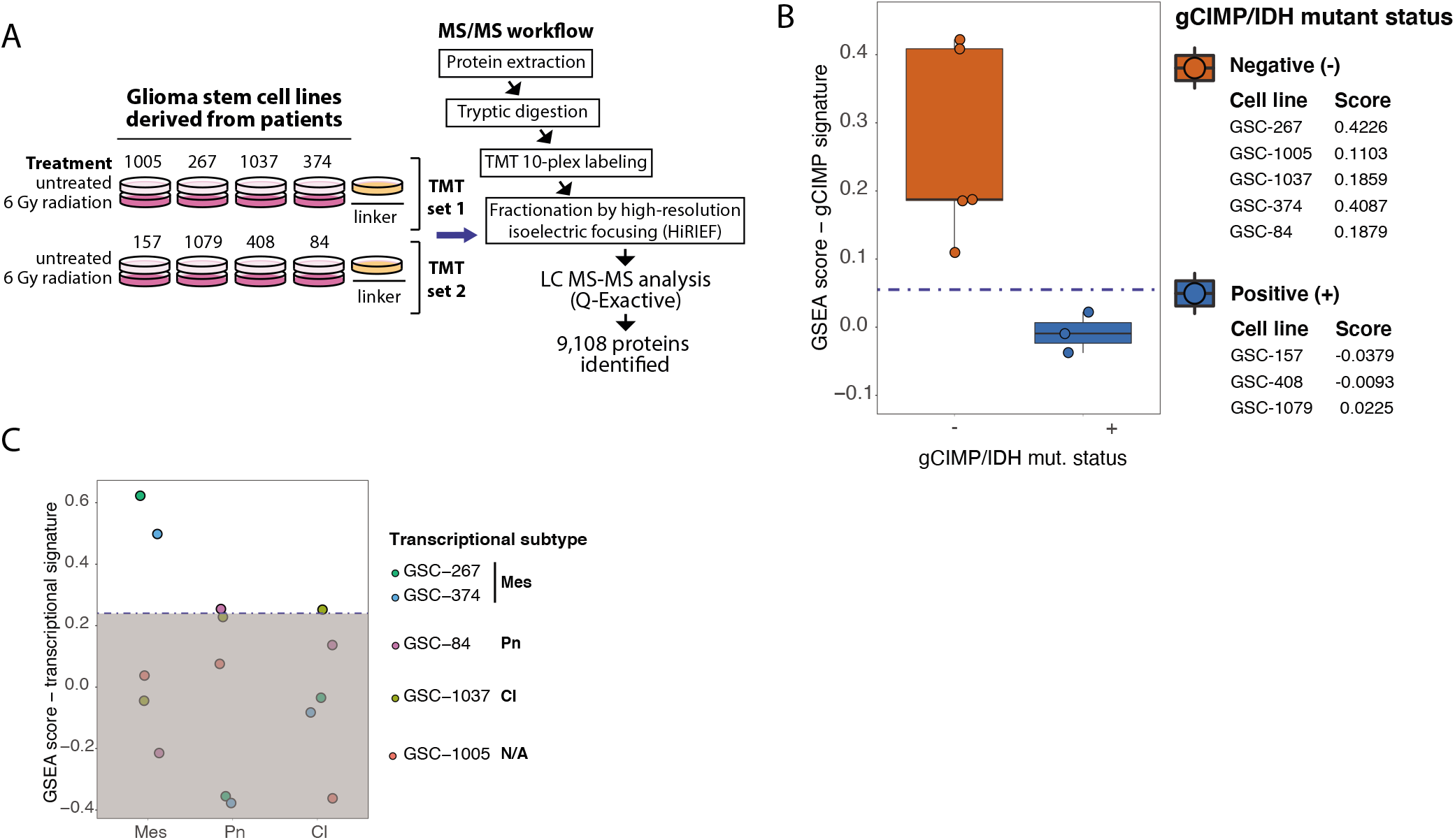
A. Workflow employed for the proteomic profiling of eight patient-derived glioma stem cell (GSC) lines. The cells were either left untreated or were treated with 6 Gray (Gy) of ionizing radiation. The number of proteins identified and quantified is displayed. TMT, Tandem Mass Tags 10plex. B. Distribution of single-sample GSEA (ssGSEA) scores representing the enrichment of a signature for cGIMP/IDH mutational status. Samples with scores that are below threshold are cGIMP/IDH mutant positive (+). C. ssGSEA scores representing the enrichment of signatures for glioma transcriptional subtypes (5). mutational status. Mes, mesenchymal; Pn, proneural; Cl, classical; N/A, ssGSEA scores below threshold for any of the subtypes.

**Supplementary Figure S2.**
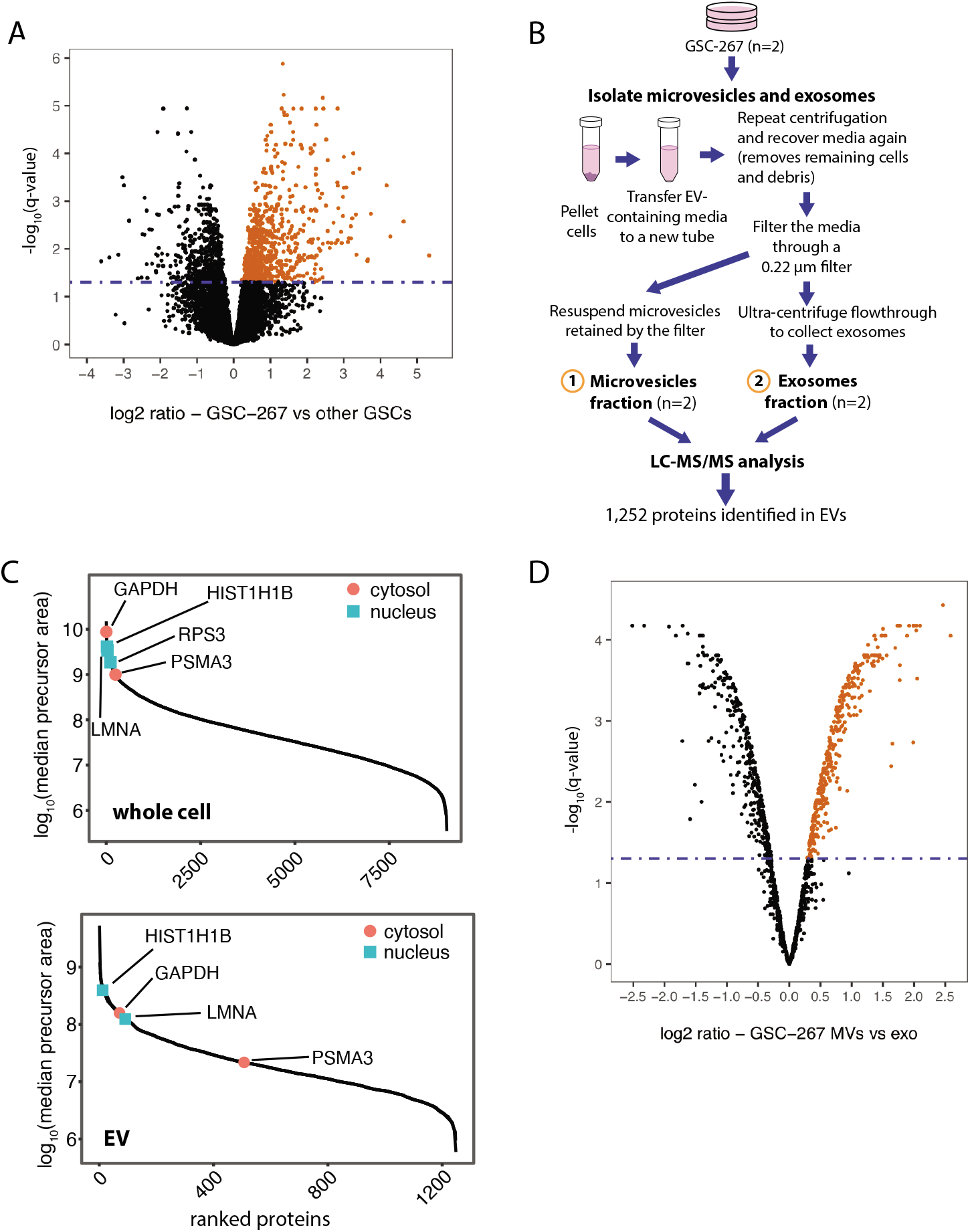
A. Volcano plots representing protein expression levels in GSC-267 cells relative to the other GSC lines plotted against the significance of the regulation based on limma statistics. Significantly elevated proteins (n = 721) are selected using the statistical package limma with a Bonferroni-Hochberg corrected q-value lower than 0.05 and log2(ratio) > 0. B. Workflow for the proteomic analysis of the cargo of EVs generated by GSCs. The approach used for isolating MVs and exosomes is outlined. Proteins are extracted from EV samples and analyzed by LC-MS. The number of proteins identified and quantified is indicated. C. Distribution of protein abundances based on precursor areas in whole cell (D) and EV (E) proteomic analyses. Cytosolic and nuclear marker proteins highlighted in red and aqua green, respectively. Marker proteins are also indicated by gene symbol. D. Volcano plots representing protein expression in MVs compared with exosomes derived from GSC-267 cells. Significantly elevated proteins (n = 356) are selected using the statistical package limma with a Bonferroni-Hochberg corrected q-value lower than 0.05 and log2(ratio) > 0.

**Supplementary Figure S3.**
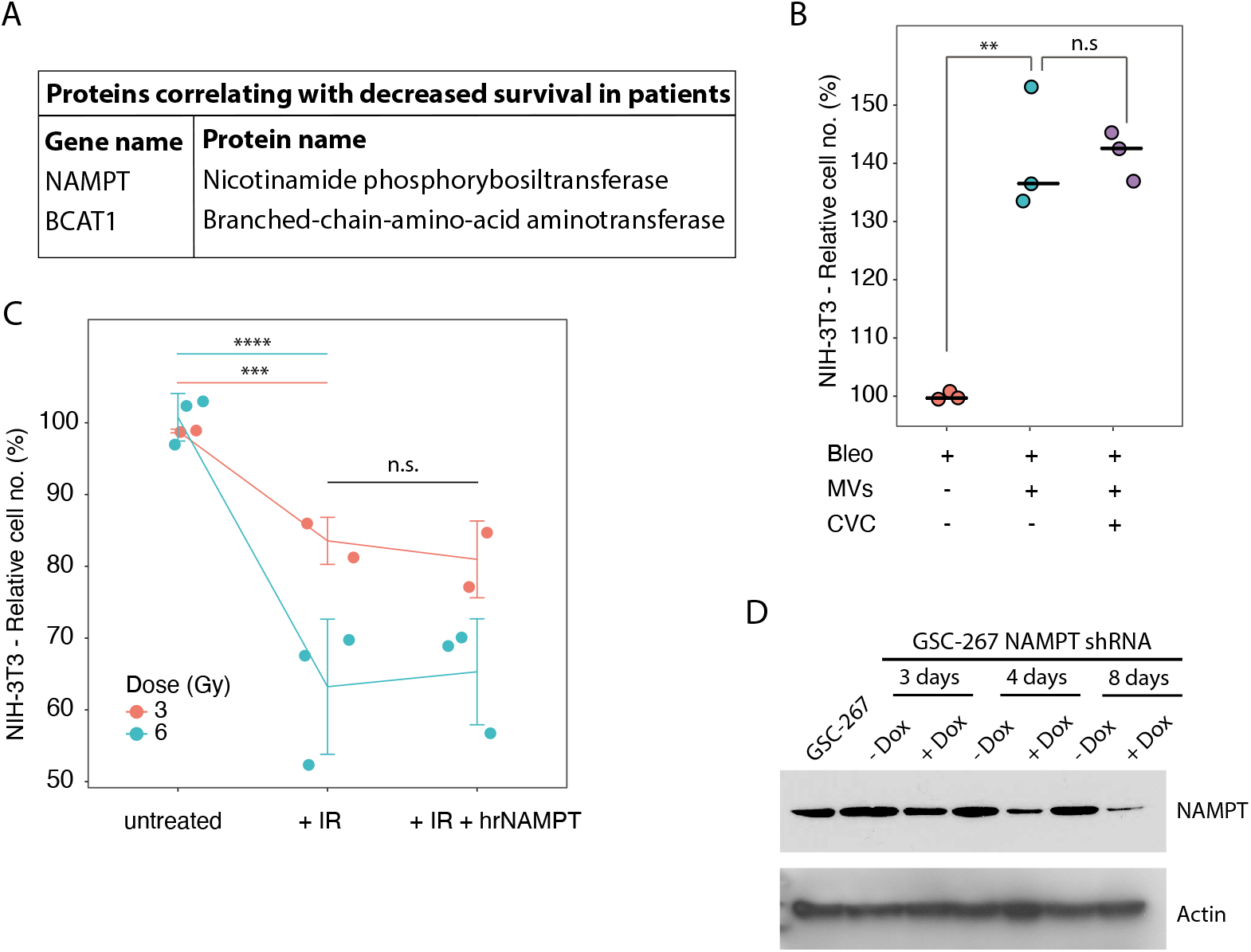
A. Table of proteins elevated in both GSC-267 cells and in the MVs that these cells produce, and which correlate with poor survival based on TCGA data. B. Relative number of viable NIH/3T3 cells cultured in low serum (0.5% CS) medium, treated with the indicated combinations of 5 μM Bleo, MVs derived from GSC-267 and 100 nM cenicriviroc (CVC) for 4 days. Individual dots represent independent biological replicates. C. Relative number of viable NIH/3T3 cells cultured in low serum (0.5% CS), either left untreated or treated with the indicated combinations of 3 or 6 Gy of ionizing radiation (IR) and 19 nM human recombinant NAMPT (hrNAMPT) for 4 days. Individual dots represent independent biological replicates. D. NAMPT protein expression levels in GSC-267 parental cells or GSC-267 NAMPTsh cells, untreated (-Dox) or treated with 400 ng/ml doxycycline (+ Dox) for the indicated length of time. Actin protein expression was used as loading control. In all figure panels, significance of observed changes was evaluated using Student’s t-test. n.s., not significant; **, p-value < 1*10^−2^; ***, p-value < 1*10^−3^; ****, p-value < 1*10^−4^.

## Notes

### Competing Interest Statement

The authors have declared no competing interest.

## References

1. S. Bao, et al., Glioma stem cells promote radioresistance by preferential activation of the DNA damage response. Nature 444, 756–760 (2006).

2. Cancer Genome Atlas Research Network, Comprehensive genomic characterization defines human glioblastoma genes and core pathways. Nature 455, 1061–8 (2008).

3. H. Yan, et al., *IDH1* and *IDH2* Mutations in Gliomas. N. Engl. J. Med. 360, 765–773 (2009).

4. H. Noushmehr, et al., Identification of a CpG Island Methylator Phenotype that Defines a Distinct Subgroup of Glioma. Cancer Cell 17, 510–522 (2010).

5. R. G. W. Verhaak, et al., Integrated Genomic Analysis Identifies Clinically Relevant Subtypes of Glioblastoma Characterized by Abnormalities in PDGFRA, IDH1, EGFR, and NF1. Cancer Cell 17, 98–110 (2010).

6. R. Stupp, et al., Radiotherapy plus Concomitant and Adjuvant Temozolomide for Glioblastoma. N. Engl. J. Med. 352, 987–996 (2005).

7. R. Stupp, et al., Maintenance therapy with tumor-Treating fields plus temozolomide vs temozolomide alone for glioblastoma a randomized clinical trial. JAMA - J. Am. Med. Assoc. 314 (2015).

8. A. Shergalis, A. Bankhead, U. Luesakul, N. Muangsin, N. Neamati, Current challenges and opportunities in treating glioblastomas. Pharmacol. Rev. 70, 412–445 (2018).

9. H. S. Friedman, et al., Bevacizumab alone and in combination with irinotecan in recurrent glioblastoma. J. Clin. Oncol. 27, 4733–40 (2009).

10. J. Y. Nam, J. F. de Groot, Treatment of Glioblastoma. https://doi.org/10.1200/JOP.2017.025536 13, 629–638 (2017).

11. J. Chen, et al., A restricted cell population propagates glioblastoma growth after chemotherapy. Nature 488, 522–526 (2012).

12. J. Skog, et al., Glioblastoma microvesicles transport RNA and proteins that promote tumour growth and provide diagnostic biomarkers. Nat. Cell Biol. 10, 1470–1476 (2008).

13. L. M. Desrochers, M. A. Antonyak, R. A. Cerione, Extracellular Vesicles: Satellites of Information Transfer in Cancer and Stem Cell Biology. Dev. Cell 37, 301–309 (2016).

14. I. Nakano, D. Garnier, M. Minata, J. Rak, Extracellular vesicles in the biology of brain tumour stem cells - Implications for inter-cellular communication, therapy and biomarker development. Semin. Cell Dev. Biol. 40, 17–26 (2015).

15. Z. Wei, et al., Coding and noncoding landscape of extracellular RNA released by human glioma stem cells. Nat. Commun. 8, 1145 (2017).

16. C. Cantó, K. J. Menzies, J. Auwerx, NAD+ Metabolism and the Control of Energy Homeostasis: A Balancing Act between Mitochondria and the Nucleus. Cell Metab. 22, 31–53 (2015).

17. A. Garten, et al., Physiological and pathophysiological roles of NAMPT and NAD metabolism. Nat. Rev. Endocrinol. 11, 535–546 (2015).

18. A. Lucena-Cacace, D. Otero-Albiol, M. P. Jiménez-García, S. Muñoz-Galvan, A. Carnero, *NAMPT* Is a Potent Oncogene in Colon Cancer Progression that Modulates Cancer Stem Cell Properties and Resistance to Therapy through Sirt1 and PARP. Clin. Cancer Res. 24, 1202–1215 (2018).

19. A. Lucena-Cacace, et al., NAMPT overexpression induces cancer stemness and defines a novel tumor signature for glioma prognosis. Oncotarget 8, 99514–99530 (2017).

20. A. D. Gujar, et al., An NAD^+^-dependent transcriptional program governs self-renewal and radiation resistance in glioblastoma. Proc. Natl. Acad. Sci. 113, E8247–E8256 (2016).

21. B. E. Kennedy, et al., NAD+ salvage pathway in cancer metabolism and therapy. Pharmacol. Res. 114, 274–283 (2016).

22. S. Lain, et al., Discovery, In Vivo Activity, and Mechanism of Action of a Small-Molecule p53 Activator. Cancer Cell 13, 454–463 (2008).

23. H. Vaziri, et al., hSIR2SIRT1 Functions as an NAD-Dependent p53 Deacetylase. Cell 107, 149–159 (2001).

24. J. Luo, et al., Negative Control of p53 by Sir2α Promotes Cell Survival under Stress. Cell 107, 137–148 (2001).

25. A. Visvanathan, et al., Essential role of METTL3-mediated m 6 A modification in glioma stem-like cells maintenance and radioresistance. Oncogene 37, 522–533 (2018).

26. A. Fidoamore, et al., Glioblastoma Stem Cells Microenvironment: The Paracrine Roles of the Niche in Drug and Radioresistance. Stem Cells Int. 2016 (2016).

27. R. Carruthers, et al., Abrogation of radioresistance in glioblastoma stem-like cells by inhibition of ATM kinase. Mol. Oncol. 9, 192–203 (2015).

28. D. Garnier, et al., Divergent evolution of temozolomide resistance in glioblastoma stem cells is reflected in extracellular vesicles and coupled with radiosensitization. Neuro. Oncol. 20, 236–248 (2018).

29. A. C. Tan, et al., Management of glioblastoma: State of the art and future directions. CA. Cancer J. Clin. 70, 299–312 (2020).

30. E. Panizza, R. M. M. Branca, P. Oliviusson, L. M. Orre, J. Lehtiö, Isoelectric point-based fractionation by HiRIEF coupled to LC-MS allows for in-depth quantitative analysis of the phosphoproteome. Sci. Rep. 7 (2017).

31. H. J. Johansson, et al., Breast cancer quantitative proteome and proteogenomic landscape. Nat. Commun. 10, 1600 (2019).

32. R. M. M. Branca, et al., HiRIEF LC-MS enables deep proteome coverage and unbiased proteogenomics. Nat. Methods 11, 59–62 (2014).

33. R. G. W. Verhaak, Moving the needle: Optimizing classification for glioma. Sci. Transl. Med. 8 (2016).

34. V. Nm, The Fundamentals of Constructing and Interpreting Heat Maps. Methods Mol. Biol. 1862, 279–291 (2019).

35. M. Fischer, Census and evaluation of p53 target genes. Oncogene 2017 3628 36, 3943–3956 (2017).

36. H. JW, A. GR, W. N, K. K, E. SJ, The p21 Cdk-interacting protein Cip1 is a potent inhibitor of G1 cyclin-dependent kinases. Cell 75, 805–816 (1993).

37. T. H, et al., A ribonucleotide reductase gene involved in a p53-dependent cell-cycle checkpoint for DNA damage. Nature 404, 42–49 (2000).

38. M. M, et al., p53 activates the CD95 (APO-1/Fas) gene in response to DNA damage by anticancer drugs. J. Exp. Med. 188, 2033–2045 (1998).

39. S. A, C. JA, J. CY, Y. JR, F. G, Bora and the kinase Aurora a cooperatively activate the kinase Plk1 and control mitotic entry. Science 320, 1655–1658 (2008).

40. Q. Wang, et al., Tumor Evolution of Glioma-Intrinsic Gene Expression Subtypes Associates with Immunological Changes in the Microenvironment. Cancer Cell 32, 42–56.e6 (2017).

41. M. Baysan, et al., G-cimp status prediction of glioblastoma samples using mRNA expression data. PLoS One 7 (2012).

42. P. Mao, et al., Mesenchymal glioma stem cells are maintained by activated glycolytic metabolism involving aldehyde dehydrogenase 1A3. Proc. Natl. Acad. Sci. U. S. A. 110, 8644–9 (2013).

43. D. Sturm, et al., Hotspot Mutations in H3F3A and IDH1 Define Distinct Epigenetic and Biological Subgroups of Glioblastoma. Cancer Cell 22, 425–437 (2012).

44. J. Behnan, G. Finocchiaro, G. Hanna, The landscape of the mesenchymal signature in brain tumours. Brain 142, 847–866 (2019).

45. L. B. Wang, et al., Proteogenomic and metabolomic characterization of human glioblastoma. Cancer Cell 39, 509–528.e20 (2021).

46. Q. Wang, et al., Tumor Evolution of Glioma-Intrinsic Gene Expression Subtypes Associates with Immunological Changes in the Microenvironment. Cancer Cell 32, 42–56.e6 (2017).

47. E. Willms, et al., Cells release subpopulations of exosomes with distinct molecular and biological properties. Sci. Rep. 6, 1–12 (2016).

48. J. DK, et al., Reassessment of Exosome Composition. Cell 177, 428–445.e18 (2019).

49. L. M. Orre, et al., SubCellBarCode: Proteome-wide Mapping of Protein Localization and Relocalization. Mol. Cell 73, 166–182.e7 (2019).

50. G. AA, T. C, G. AA, S. JK, Extracellular nicotinamide phosphoribosyltransferase, a new cancer metabokine. Br. J. Pharmacol. 173, 2182–2194 (2016).

51. T. S, et al., The Cytokine Nicotinamide Phosphoribosyltransferase (eNAMPT; PBEF; Visfatin) Acts as a Natural Antagonist of C-C Chemokine Receptor Type 5 (CCR5). Cells 9 (2020).

52. R. D, et al., Macrophages provide a transient muscle stem cell niche via NAMPT secretion. Nature 591, 281–287 (2021).

53. X. Jiao, et al., CCR5 governs DNA damage repair and breast cancer stem cell expansion. Cancer Res. 78, 1657 (2018).

54. Q. Feng, et al., A class of extracellular vesicles from breast cancer cells activates VEGF receptors and tumour angiogenesis. Nat. Commun. 8 (2017).

55. L. M. Desrochers, F. Bordeleau, C. A. Reinhart-King, R. A. Cerione, M. A. Antonyak, Microvesicles provide a mechanism for intercellular communication by embryonic stem cells during embryo implantation. Nat. Commun. (2016) https://doi.org/10.1038/ncomms11958.

56. M. Yoshida, et al., Extracellular Vesicle-Contained eNAMPT Delays Aging and Extends Lifespan in Mice. Cell Metab. 30, 329–342.e5 (2019).

57. E. Van Der Veer, et al., Extension of human cell lifespan by nicotinamide phosphoribosyltransferase. J. Biol. Chem. 282, 10841–10845 (2007).

58. S. Imai, J. Yoshino, The importance of NAMPT/NAD/SIRT1 in the systemic regulation of metabolism and ageing. Diabetes, Obes. Metab. 15, 26–33 (2013).

59. Y. K, O. K, H. K, N. T, NAD Metabolism in Cancer Therapeutics. Front. Oncol. 8 (2018).

60. S. D, Z. TS, M. DL, O. T, D. PS, Inhibition of nicotinamide phosphoribosyltransferase (NAMPT) as a therapeutic strategy in cancer. Pharmacol. Ther. 151, 16–31 (2015).

61. P. G. Moon, et al., Identification of developmental endothelial locus-1 on circulating extracellular vesicles as a novel biomarker for early breast cancer detection. Clin. Cancer Res. 22, 1757–1766 (2016).

62. A. A. Grolla, et al., Nicotinamide phosphoribosyltransferase (NAMPT/PBEF/visfatin) is a tumoural cytokine released from melanoma. Pigment Cell Melanoma Res. 28, 718–729 (2015).

63. S. Rontogianni, et al., Proteomic profiling of extracellular vesicles allows for human breast cancer subtyping. Commun. Biol. 2, 1–13 (2019).

64. X. Shi, et al., NAD+ depletion radiosensitizes 2-DG-treated glioma cells by abolishing metabolic adaptation. Free Radic. Biol. Med. 162, 514–522 (2021).

65. K. P. L. Bhat, et al., Mesenchymal differentiation mediated by NF-κB promotes radiation resistance in glioblastoma. Cancer Cell 24, 331–46 (2013).

66. H. Jeong, et al., Correcting for Naturally Occurring Mass Isotopologue Abundances in Stable-Isotope Tracing Experiments with PolyMID. Metab. 2021, Vol. 11, Page 310 11, 310 (2021).

67. D. J. Clark, et al., Integrated Proteogenomic Characterization of Clear Cell Renal Cell Carcinoma. Cell 179, 964–983.e31 (2019).

68. Y. Dou, et al., Proteogenomic Characterization of Endometrial Carcinoma. Cell 180, 729–748.e26 (2020).

69. C. Huang, et al., Proteogenomic insights into the biology and treatment of HPV-negative head and neck squamous cell carcinoma. Cancer Cell 39, 361–379.e16 (2021).

70. Y. Hu, et al., Integrated Proteomic and Glycoproteomic Characterization of Human High-Grade Serous Ovarian Carcinoma. Cell Rep. 33 (2020).

71. Y. H. Hur, R. A. Cerione, M. A. Antonyak, Extracellular vesicles and their roles in stem cell biology. Stem Cells 38, 469–476 (2020).

72. A. Latifkar, Y. H. Hur, J. C. Sanchez, R. A. Cerione, M. A. Antonyak, New insights into extracellular vesicle biogenesis and function. J. Cell Sci. 132 (2019).

73. E. Panizza, R. A. Cerione, M. A. Antonyak, Exosomes as Sentinels against Bacterial Pathogens. Dev. Cell (2020) https://doi.org/10.1016/j.devcel.2020.03.022.

74. W. F, C. RA, A. MA, Isolation and characterization of extracellular vesicles produced by cell lines. STAR Protoc. 2, 100295 (2021).

75. P. Hovstadius, A Phase I study of CHS 828 in patients with solid tumor malignancy. Clin. Cancer Res. 8, 2843–2850 (2002).

76. A. Ravaud, Phase I study and pharmacokinetic of CHS-828, a guanidino-containing compound, administered orally as a single dose every 3 weeks in solid tumours: an ECSG/EORTC study. Eur. J. Cancer 41, 702–707 (2005).

77. S. Goldinger, Efficacy and safety of APO866 in patients with refractory or relapsed cutaneous T-cell lymphoma: a phase 2 clinical trial. JAMA Dermatol. 152, 837–839 (2016).

78. M. Pishvaian, A phase I trial of GMX1777, an inhibitor of nicotinamide phosphoribosyl transferase (NAMPRT), given as a 24-hour infusion. J. Clin. Oncol. 27, 3581–3581 (2009).

79. L. S. E. H. K. B. A. H. K Holen, The pharmacokinetics, toxicities, and biologic effects of FK866, a nicotinamide adenine dinucleotide biosynthesis inhibitor. Invest. N. Drugs 26, 45–51 (2008).

80. A. B. R. L. P. N. A Heideman, Safety and efficacy of NAD depleting cancer drugs: results of a phase I clinical trial of CHS 828 and overview of published data. Cancer Chemother. Pharmacol. 65, 1165–1172 (2010).

81. Y. Zhu, et al., Exogenous NAD+ decreases oxidative stress and protects H2O2-treated RPE cells against necrotic death through the up-regulation of autophagy. Sci. Rep. 6 (2016).

82. E. Dolgin, Anticancer autophagy inhibitors attract “resurgent” interest. Nat. Rev. Drug Discov. 18, 408–410 (2019).

83. M. Rangel, J. Kong, V. Bhatt, K. Khayati, J. Y. Guo, Autophagy and tumorigenesis. FEBS J. (2021) https://doi.org/10.1111/FEBS.16125.

84. M. Hallschmid, H. Randeva, B. K. Tan, W. Kern, H. Lehnert, Relationship Between Cerebrospinal Fluid Visfatin (PBEF/Nampt) Levels and Adiposity in Humans. Diabetes 58, 637–640 (2009).

85. N. Hara, K. Yamada, T. Shibata, H. Osago, M. Tsuchiya, Nicotinamide Phosphoribosyltransferase/Visfatin Does Not Catalyze Nicotinamide Mononucleotide Formation in Blood Plasma. PLoS One 6 (2011).

86. J. L. Welton, et al., Cerebrospinal fluid extracellular vesicle enrichment for protein biomarker discovery in neurological disease; multiple sclerosis. J. Extracell. vesicles 6 (2017).

87. C. Macron, et al., Exploration of human cerebrospinal fluid: A large proteome dataset revealed by trapped ion mobility time-of-flight mass spectrometry. Data Br. 31, 105704 (2020).

88. V. Audrito, et al., Extracellular nicotinamide phosphoribosyltransferase (NAMPT) promotes M2 macrophage polarization in chronic lymphocytic leukemia. Blood 125, 111–123 (2015).

89. C. Gu, et al., Tumor-specific activation of the C-JUN/MELK pathway regulates glioma stem cell growth in a p53-dependent manner. Stem Cells 31, 870–881 (2013).

90. C. S. Hughes, et al., Ultrasensitive proteome analysis using paramagnetic bead technology. Mol. Syst. Biol. 10, 757 (2014).

91. S. M, et al., Evaluation of FASP, SP3, and iST Protocols for Proteomic Sample Preparation in the Low Microgram Range. J. Proteome Res. 16 (2017).

92. J. D. Holman, D. L. Tabb, P. Mallick, Employing ProteoWizard to Convert Raw Mass Spectrometry Data. Curr. Protoc. Bioinforma. 46, 13.24.1–13.24.9 (2014).

93. V. Granholm, et al., Fast and accurate database searches with MS-GF+Percolator. J. Proteome Res. 13, 890–897 (2014).

94. J. Boekel, et al., Multi-omic data analysis using Galaxy. Nat. Biotechnol. 2015 332 33, 137–139 (2015).

95. M. Sturm, et al., OpenMS - An open-source software framework for mass spectrometry. BMC Bioinformatics 9, 1–11 (2008).

96. S. Hänzelmann, R. Castelo, J. Guinney, GSVA: Gene set variation analysis for microarray and RNA-Seq data. BMC Bioinformatics 14 (2013).

97. M. E. Ritchie, et al., limma powers differential expression analyses for RNA-sequencing and microarray studies. Nucleic Acids Res. 43, e47–e47 (2015).

98. A. A, R. J, topGO: Enrichment Analysis for Gene Ontology. (2021) https://doi.org/10.18129/B9.bioc.topGO.

99. D. Szklarczyk, et al., The STRING database in 2017: quality-controlled protein-protein association networks, made broadly accessible. Nucleic Acids Res. 45, D362–D368 (2017).

100. M. Pernemalm, et al., In-depth human plasma proteome analysis captures tissue proteins and transfer of protein variants across the placenta. Elife 8 (2019).

101. B. CW, et al., The somatic genomic landscape of glioblastoma. Cell 155, 462 (2013).

102. J. E. Blum, et al., Pyruvate Kinase M2 Supports Muscle Progenitor Cell Proliferation but Is Dispensable for Skeletal Muscle Regeneration after Injury. J. Nutr. 151, 3313–3328 (2021).

103. C. A. Schneider, W. S. Rasband, K. W. Eliceiri, NIH Image to ImageJ: 25 years of image analysis. Nat. Methods 2012 97 9, 671–675 (2012).

104. B. T. Kreger, E. R. Johansen, R. A. Cerione, M. A. Antonyak, The Enrichment of Survivin in Exosomes from Breast Cancer Cells Treated with Paclitaxel Promotes Cell Survival and Chemoresistance. Cancers 2016, Vol. 8, Page 111 8, 111 (2016).

